# Alternative splicing and its regulation in the malaria vector *Anopheles gambiae*

**DOI:** 10.1101/2023.07.18.549290

**Authors:** Bárbara Díaz-Terenti, Elena Gómez-Díaz

## Abstract

Alternative splicing (AS) is a highly conserved mechanism that allows to expand the coding capacity of the genome, by modifying how multiple isoforms are expressed or used to generate different phenotypes. Despite its importance in physiology and disease, genome-wide studies of AS are lacking in most insects, including mosquitoes. Even for model organisms, chromatin associated processes involved in the regulation AS are poorly known. In this study, we investigated AS in the mosquito *Anopheles gambiae* in the context of tissue-specific gene expression and mosquito responses to a *Plasmodium falciparum* infection, as well as the relationship between patterns of differential isoform expression and usage with chromatin accessibility changes. For this, we combined RNA-seq and ATAC-seq data from *A. gambiae* midguts and salivary glands, and from infected and non-infected midguts. We report differences between tissues in the expression of 456 isoforms and in the use of 211 isoforms. Secondly, we find a clear and significant association between chromatin accessibility states and tissue-specific patterns of AS. The analysis of differential accessible regions located at splicing sites permitted the identification of several motifs resembling the binding sites of *Drosophila* transcription factors. Finally, the genome-wide analysis of tissue-dependent enhancer activity revealed that approximately 20% of *A. gambiae* transcriptional enhancers annotate to a differentially expressed or used isoform and that their activation status is linked to AS differences between tissues. This research illuminates the role of AS in gene expression in vector mosquitoes, and identifies regulatory regions potentially involved in AS regulation, which could reveal novel strategies for vector control.

## 1 Introduction

Alternative splicing (AS from here) is an important source of phenotypic variation in eukaryotes that allows the generation of multiple transcripts from a single gene, expanding the coding capacity and functionality of the genome. AS regulates gene expression by acting on mRNA transcripts at a post-transcriptional level and is critical for diverse cellular processes, including cell differentiation and development as well as cell reprogramming and tissue remodeling.

AS is mediated by four types of basic events: alternative 5′ splice-site, alternative 3′ splice-site, cassette-exon inclusion or skipping, and intron retention. These processes can be combined to obtain dozens of isoforms from a single gene. On the other hand, AS regulation include different mechanisms as the temporal and spatial differential expression of the isoforms, the activation of the RNA Pol II or the on-off regulation by nonsense-mediated decay (Wang and Cooper, 2007, Nilsen and Graveley, 2010, Kelemen et al., 2013, Shenasa and Hertel, 2019, Ule and Blencowe, 2019, Wright, Smith and Jiggins, 2022). Another process associated to the AS is regulating isoform usage. Isoform usage is defined as, the participation of each isoform in the gene expression, so when these proportions switch between two conditions, some isoforms are differentially used (Vitting-Seerup and Sandelin, 2017, Merino, Conesa and Fernández, 2019, De La Fuente et al., 2020, Li et al., 2020a, Chen et al., 2022). As a result, the function of gene might change because different variant proteins are produced, while the total gene expression can remain unaltered.

The type of events and mechanisms of AS vary significantly amongst species. Despite of this, AS is highly conserved in eukaryotes playing fundamental functions in health and disease, adaptation to the environment, and defence against pathogens (Nilsen and Graveley, 2010, Kelemen et al., 2013, Martinez and Lynch, 2013, Bush et al., 2017, Vitting-Seerup and Sandelin, 2017, Ule and Blencowe, 2019, Zhao, 2019, Li et al., 2020a, Song et al., 2020, Verta and Jacobs, 2022, Wright, Smith and Jiggins, 2022).

A simple way to study AS is to quantify and compare the expression of isoforms that are generated in different tissues or experimental conditions (Katz et al., 2010, Glaus, Honkela and Rattray, 2012). Transcriptomic data obtained by RNA-seq is mostly used for gene level analysis (Vitting-Seerup and Sandelin, 2017, Zhao, 2019), but offers the possibility to study AS to analyze and count known transcripts or to detect new splicing events (Zhao, 2019).

Regulation of AS is intrinsically related to chromatin dynamics. To obtain diverse transcripts, *cis* and *trans*-regulatory elements participate both in the generation of transcripts and in marking splice sites (Bush et al., 2017, Shenasa and Hertel, 2019, Zhao, 2019, Verta and Jacobs, 2022). In addition, AS regulation can be affected by posttranslational histone modifications, nucleosome positioning, and chromatin accessibility (Davuluri et al., 2008, Luco et al., 2011, Montes et al., 2012, Klemm, Shipony and Greenleaf, 2019, Nussinov et al., 2021). However, how these different epigenetic layers interact, and which are the mechanisms that underlie the great diversification of gene functions through AS, has been poorly investigated (Davuluri et al., 2008, Luco et al., 2011, Klemm, Shipony and Greenleaf, 2019). The integrated analysis of transcriptomic and epigenetic data could bring new insights into how AS is regulated and how it changes under different conditions.

AS has been widely studied in model organisms like *Drosophila melanogaster*, *Danio renio* or *Caenorhabditis elegans* (Zahler, 2005, Venables, Tazi and Juge, 2012, Lin et al., 2022). In *Drosophila melanogaster* AS plays important roles in sex-determination, muscle, neural development among other biological processes (Venables, Tazi and Juge, 2012). More precisely the relationship between AS and chromatin has been investigated in *Drosophila* gonads. That study revealed that the splicing factors expressed in the undifferentiated cells act coordinately with chromatin remodeling factors and histone modifying enzymes to maintain the transcription network through post-transcriptional mechanisms (Gan et al., 2010). Despite of this, there is a lack of the knowledge in how the AS interacts with chromatin to regulate gene function under different physiological or cellular conditions (Nilsen and Graveley, 2010, Luco et al., 2011, Kelemen et al., 2013).

In mosquitoes, little is known about AS and its regulation. Given that AS is evolutionarily conserved, one would expect that functions and mechanisms are maintained across insects. There is, however, marked inter-species variation in alternative splicing that probably resulted through selective pressures exerted by environment and lifestyle (Kurtz and Armitage, 2006, Malko et al., 2006). In *Aedes aegypti*, previous studies have focused on sex differentiation (Salvemini et al., 2011, 2013, Calkins et al., 2015). In the human malaria mosquito *Anopheles gambiae*, the study of AS is limited to a few genes, such as *doublesex* and *Dscam* (Tsujimoto et al., 2013, Krzywinska et al., 2016, Sinkins, 2016, Djihinto et al., 2022), tissues like salivary glands (Dixit et al., 2009) or signalling pathways (Meister et al., 2005). These studies suggest that AS could play a crucial role in vector competence and mosquito immunity. However, a more systematic analysis onto the role of AS and its regulation in mosquitoes, is lacking.

In this study, we combined transcriptomic and chromatin accessibility data of infected and non-infected *A. gambiae* salivary glands and midguts to interrogate about AS regulation. In particular, which isoforms belonging to multisoform genes are differentially expressed and used in different mosquito tissues infected with the malaria parasite *Plasmodium falciparum*, and whether chromatin-associated mechanisms underlie these events (Love, Huber and Anders, 2014, Vitting-Seerup, Sandelin and Berger, 2019). We contend that a deeper understanding of the mechanisms of AS regulation can reveal key *cis* and *trans* regulatory elements, like splicing transcription factors and enhancers, involved in mosquito responses to *P. falciparum* infection and/or vector competence.

## 2 Results

### 2.1 Alternative splicing (AS) regulates tissue-specific gene expression in infected *A. gambiae*

Considering the 1,263 multisoform genes detected in *A. gambiae*, we aimed to identify alternative splicing events and their role in gene expression regulation in different tissues and infection conditions. For this, we performed two types of analysis: one to identify differentially expressed isoforms belonging to multisoform genes (DEMG), and another to detect isoforms differentially used (DUI); isoforms that are altered in their relative proportion to the total expression of the gene.

In the comparison between *P. falciparum* infected and non-infected (control) mosquito midguts, 32 isoforms appeared differentially regulated out of the 848 multisoform genes present in this dataset (with counts > 10 as the detectability threshold). Of these, 19 (2%) correspond to differentially expressed and 13 (1.5%) to differentially used isoforms. In salivary glands, we detected 977 expressed multisoform genes of which 91 appear differentially regulated; 56 (6%) were differentially expressed and 35 (3.5%) differentially used.

Then, we examined differences in AS regulation between tissues: midguts and salivary glands infected with *P. falciparum*. We detect 920 multisoform genes: 456 (50%) DEMG and 211 (23%) DUI.

We also report multisoform genes in which, one or more isoforms were differentially expressed and used (Fig 1a). In our results, the overlap between AS mechanisms represented 22% (105) of the multisoform genes that display AS when comparing midguts and salivary glands (Fig 1b), and less than 1% (0 for the midguts and 1 for the salivary glands) when comparing infected and non-infected mosquito tissues (Supplementary table S1 and S2).

**Fig 1.**
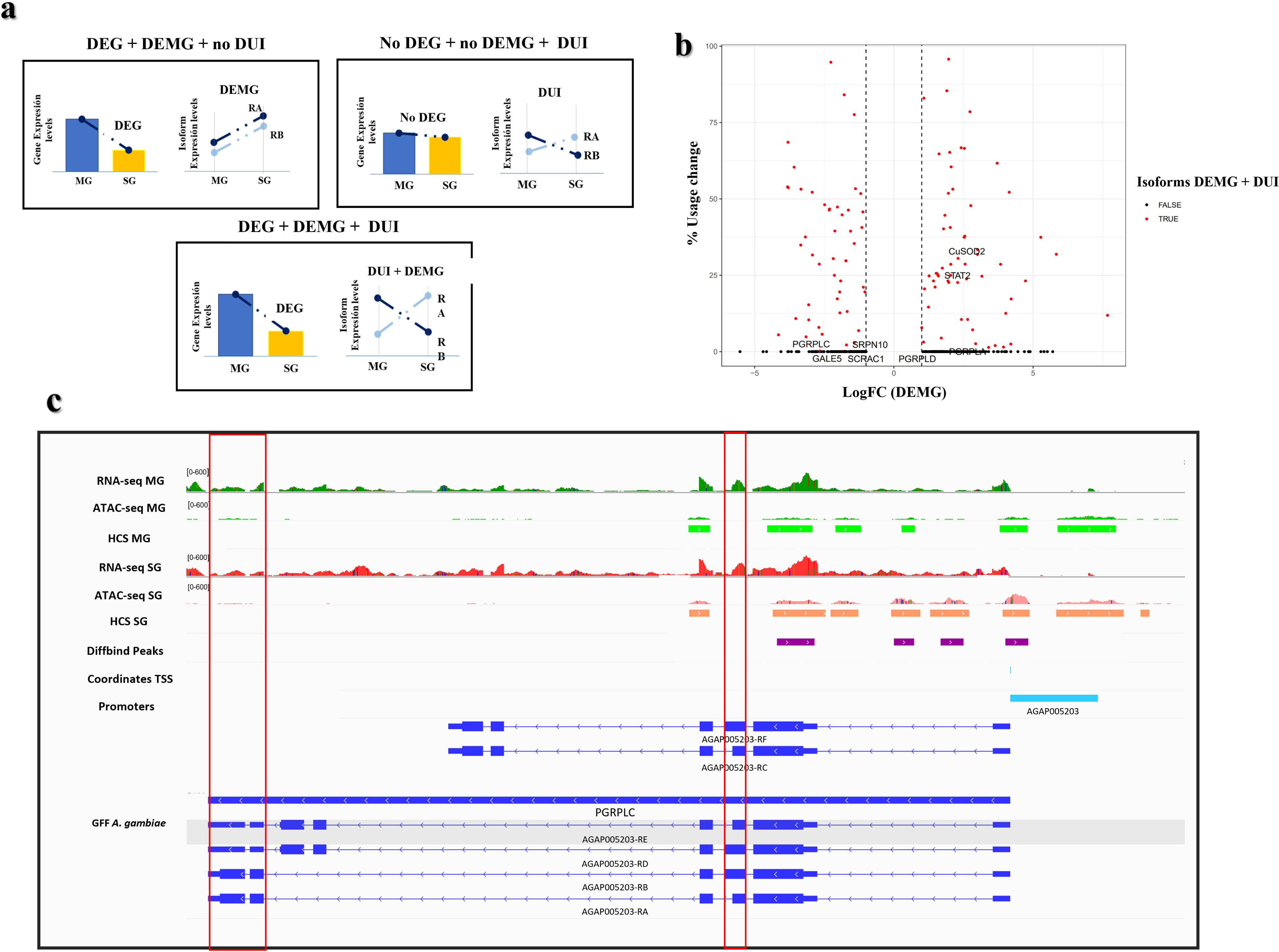
Patterns of differentially expressed isoforms belonging to multisoform genes (DEMG) and differentially used isoforms between tissues. **a**, Diagram illustrating possible DEMG and DUI patterns expected in our data. The gene can be differentially expressed, and the isoform differentially expressed but not differentially used, only one more of the isoforms differentially used and the isoforms can be differentially expressed and used at the same time; **b**, Volcano plot showing total usage change (i.e., expression redistribution between isoforms) versus log-transformed values of isoform expression fold change between tissues. Isoforms that are DUI and DEMG are shown in red whereas black dots correspond to differentially expressed isoforms. Labels identify isoforms corresponding to *A. gambiae* genes with immune functions; **c**, IGV representation of the AGAP005203 (PGRPLC) gene. Red lines highlight differential expression and accessibility at RA and RE differential exons. The red box on the left points to a higher expression of the RA isoform in midguts, while the box on the right points to a higher expression in salivary glands.

To test for differential functionality of DEMG and DUI isoforms between tissues, we performed GO enrichment analysis. For the differentially expressed isoforms that appear more expressed in midguts, the 10 first GO terms (ordered by less p-value) include metabolic process, regulation of response to stimulus and signaling. Contrary, differentially expressed isoform genes that are more expressed in salivary glands correspond to functions related to fiber and cytoskeleton organization, ion transport, and muscle contraction (Supplementary table S3).

In the case of differentially used isoforms that appear overused in midguts, the 10 first GO terms correspond to functions related to muscle, contraction, development and differentiation, proton transmembrane transport and metabolic processes. In salivary glands, overused isoforms displayed functions like metabolic process of nucleotides and carboxylic acids, and muscle contraction (Supplementary table S3).

Next, we described the types of events that lead to differential isoform expression (DEMG) and/or isoform usage (DUI) in our set of *A.gambiae* multisoform genes. Our results show that Exon Skipping (ES) and Alternative Start Site (ATSS) are the two most frequent types of events. For differentially expressed isoforms, we report differences in the frequency of these events between conditions for both ES and ATSS: 68 ES events in *Inf MG vs.* 102 ES events in *Inf SG* and in the case of ATSS, 89 ATSS events in *Inf MG vs.* 105 ATSS events in *Inf SG.* For the differentially used isoforms, only the frequency of ATSS is moderately altered when comparing Inf MG vs. Inf SG (*43 ATSS events in SG compared to 51 events in MG*).

Finally, we interrogated whether our differentially expressed and used isoforms coincided with genes known to be involved in mosquito immunity (Table S4) (Sreenivasamurthy et al., 2013). We found 8 genes among our differentially expressed and used isoforms, that encode for proteins involved in anti-*Plasmodium* mosquito responses (STAT, PGRPLC, OXR1). One example are the AGAP005203-RA and AGAP005203-RE isoforms of the PGRPLC gene, identified in the present work as differentially expressed (DEMG). The RA isoform is overexpressed during midgut infection, while RE isoform, is overrepresented in infected salivary glands compared to control. RB and RC appear similarly expressed, and the rest of isoforms have values below the expression threshold, so they were considered as non-expressed (Fig 1c).

### 2.2 Alternative Splicing is positively correlated with chromatin accessibility in malaria-infected *A. gambiae* tissues

#### 2.2.1 Tissue-specific isoform expression is positively associated to differential chromatin accessibility

First, we tested if higher expression of the isoforms was associated with higher levels of chromatin accessibility in the isoform promoter (1kb upstream of the TSS/ATG of the isoform) or the isoform body. The results for the differentially expressed group show a positive and significant correlation between isoform expression and chromatin accessibility for the isoform body, both in salivary glands (*rho* = 0,328, *p-value* = 8,095e-07) and midguts (*rho* = 0,369, *p-value* = 3.009e-07), but not for the promoter region (Table S5A) (Fig 2).

**Fig 2.**
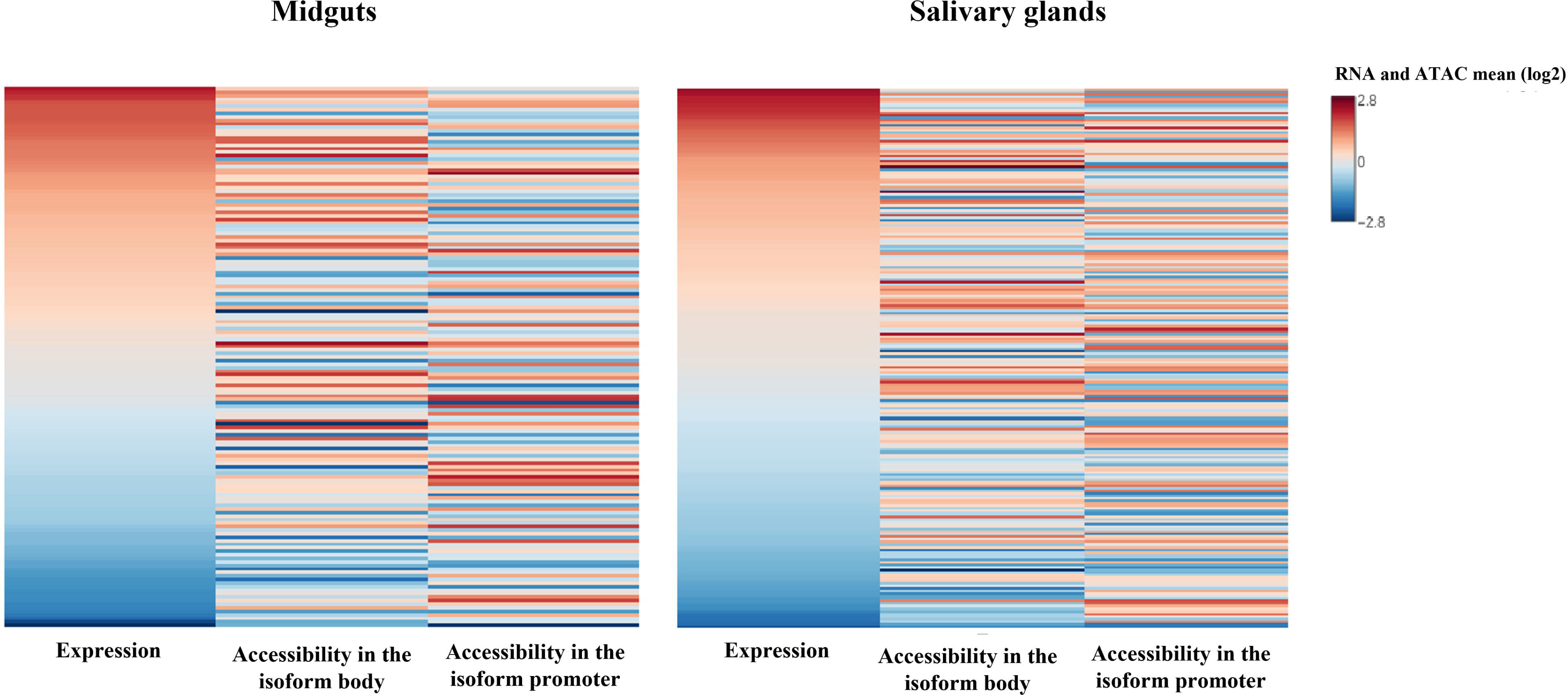
Differential expression of the isoforms correlates with changes in chromatin accessibility between tissues in the isoform body. Heatmap showing ATAC-seq nucleosome-free enrichment at promoters or in the body of DEMG isoforms and their expression levels. Genes are ordered by normalized RNA-seq isoform expression (RPKM). Changes tend to occur in the same direction, and there is a positive and significant correlation between the magnitude of changes in accessibility and expression at the isoform body in both tissues.

Next, we examined whether the relationship is quantitative, that is, if the levels of expression of the isoform differentially expressed (categorized as high, medium, or low) were correlated with the degree of chromatin accessibility in the isoform promoter. This relationship appears not significant either in midguts or salivary glands (*rho* < 0,02, *p-value* > 0.1 for all levels and tissues, Table S5B) (Fig S2).

For the analysis of chromatin accessibility in relation to differential usage, we used processed data of differential accessibility peaks previously published by us (Ruiz, Ranford-Cartwright and Gómez-Díaz, 2021). We then selected DiffBind peaks that were located at the splicing site or in the region nearby (+/-100 Kb), and that annotated to differentially used isoforms. We report a clear association between differential usage and accessibility with 98 differentially used isoforms, 54,4% of the total number of isoforms differentially used, displaying a DiffBind peak located at or nearby splicing sites.

#### 2.2.2 Motif analysis

We performed a motif analysis using HOMER on the set of DiffBind peaks that annotated to differentially used isoforms and located at or nearby splicing sites. The purpose of this analysis was to detect overrepresented motifs that point to transcription factors involved in mosquito AS regulation.

Using the hierarchical clustering of MEME *de novo motif discovery* option, several motif hits with distinct similarities with *Drosophila* sequences appear overrepresented (summary in Table 1) (Table S6B). For the subset of differentially used isoforms, we found *de novo* motifs related with brachyentron (byn), implicated in midgut development, snail (sna), which participates in epithelial transition, grainy head (grh) involved in epithelial cell fate or chromatin remodellers like the Chromatin-linked adaptor for MSL proteins (CLAMP).

Now analyzing the set of “known” motifs, we found only two motifs overrepresented. The motif CATCMCTA matches a *Drosophila* promoter element of unknown function, and the motif GGYCATAAAW corresponds to the caudal homeobox of *Drosophila* (Table S6A).

### 2.3 Mosquito enhancers involved in AS

We aimed to study if *A. gambiae* previously identified transcriptional enhancers which participated in the regulation of AS. Such a relationship has been shown in human and mouse in which 5% of the enhancers appear associated to AS changes (Shiau et al., 2021). For this analysis, we used *A. gambiae* enhancers described by our lab in a previous study (Ruiz, Ranford-Cartwright and Gómez-Díaz, 2021). This dataset included 4,268 enhancers identified computationally by orthology with *Drosophila* enhancers, or experimentally discovered by others in mosquitoes, and that were validated by ours using the combined analysis of differential chromatin accessibility and histone modifications marking, as well as gene expression changes between tissues (Ruiz, Ranford-Cartwright and Gómez-Díaz, 2021). Of these mosquito enhancers, 811 appear active in our dataset.

In the differentially expressed isoforms we found a total of 125 accessible enhancers that annotate to 76 genes (21,1% of the 360 differentially expressed isoforms belonging to multisoform genes) (Supplementary Table S7A). Of these 125 accessible (active) enhancers, 26 of them coincide with a Diffbind peak, that is, they are differentially accessible between infected midguts and salivary glands.

For the group of differentially used isoforms, we identified 39 accessible enhancer regions that annotate to 23 genes (19,8% of the 116 differentially used isoforms belonging to multisoform genes) (Supplementary Table S7B), and 11 exhibit Diffbind peaks, indicating differential accessibility between infected midguts and salivary glands.

Lastly, we examined the location of the enhancers annotated to differentially expressed and used isoforms. For both, the DEMG and DUI groups, the number of intragenic enhancers almost double of the number of intergenic (DEMG: 44 intergenic and 81 intragenic enhancers; DUI: 14 intergenic and 25 intragenic enhancers).

One example is the gene AGAP005234 that encodes for a copper-zinc superoxide dismutase. This protein has functions involved in mosquito immunity, including anti-Plasmodium responses, as well as response to oxygen radicals. (Marikovsky et al., 2003, Molina-Cruz et al., 2008). The gene contains two different isoforms (AGAP005234-RA and AGAP005234-RB), RB isoform is differentially expressed and used in salivary glands and RA is differentially used in midguts. The enhancer associated to this gene is intergenic, located 2 Kbp upstream of the start of the gene, and its accessibility in higher in SG coinciding with the expression and use that is also higher in SG (Fig 3).

**Fig 3.**
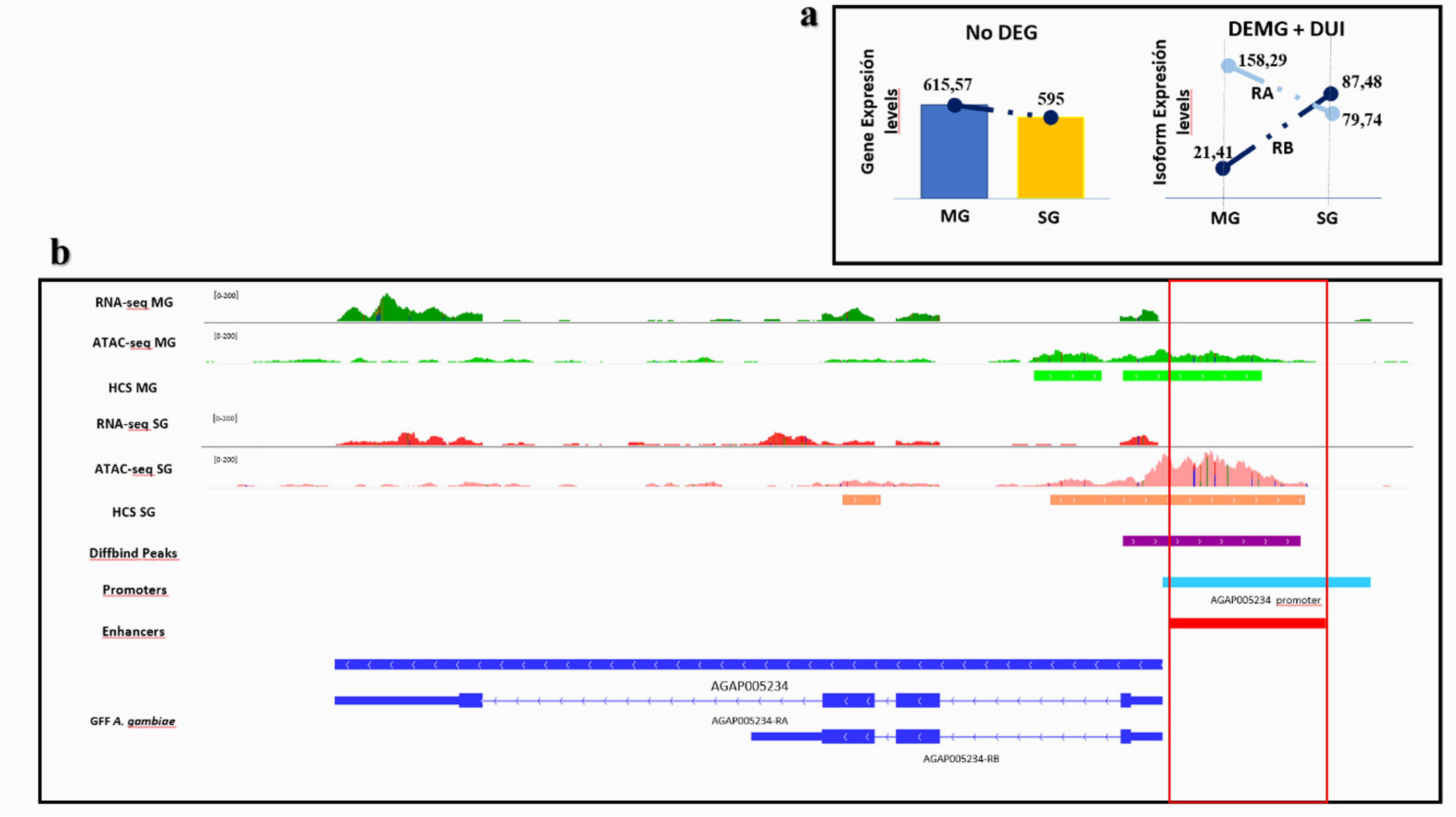
Enhancer function and alternative splicing of the gene AGAP005234. **a**, Scheme summarizing gene and isoform expression between tissues; **b**, IGV screenshot showing the enhancer annotated to the AGAP005234 gene which is differentially accessible between tissues coinciding with the differential expression and usage of their isoforms. RB isoform is differentially expressed and used in salivary glands and the enhancer is more accessible in this tissue. All tracks are shown at equal scale.

## 3 Discussion

Alternative splicing has been relatively little studied in mosquitoes, and the relationship of this process with chromatin dynamics remains mostly unknown. In this study we investigated alternative splicing in *A. gambiae* and the role of chromatin structure in its regulation. We compared patterns of AS in different conditions, first between two mosquito tissues: midguts and salivary glands, and second between *P. falciparum* infected and non-infected (control) mosquito midguts and salivary glands.

The first part of this work aimed at identifying how prevalent is AS in mosquitoes and how AS patterns change between tissues and infection status. From a total of 1,263 *A. gambiae* genes susceptible of AS, we detect around a 67-77% being differentially expressed or used at the level of isoforms.

These differences were largely due AS events at the tissue-level. Indeed, when comparing the two infected tissues, the 49% of isoforms (456) appeared differentially expressed (DEMG) and 22.9% (211) differentially used (DUI), suggesting that AS plays an important role in regulating tissue-specific expression in *A. gambiae*. The role of AS in tissue specific processes has been reported in different organisms like in humans, *D. melanogaster*, *C. elegans* (Wang et al., 2008, Telonis-Scott et al., 2009, Venables, Tazi and Juge, 2012, Ragle et al., 2015) or even in *Anopheles stephensi* (Sreenivasamurthy et al., 2017).

However, while in this study the AS patterns vary significantly between tissues, differences were minimal when comparing infected and uninfected tissues for both groups, differentially expressed, and used isoforms (only 32 DEMG and DUI isoforms detected in case of midgut infection and 91 DEMG and DUI isoforms detected in case of salivary glands infection). These results can be explained by the inter-sample variability in these comparisons as revealed the PCA analysis. That may be reducing the statistical power of the expression and differential use tests, resulting in low numbers of differentially expressed and used isoforms.

Among the tissue-specific multisoform genes, we report several genes with immune functions that have a specific isoform differentially expressed in midguts and another isoform differentially expressed in salivary glands, meaning that each has a different function depending on the tissue in which it is expressed. An example of this is the AGAP005203 gene, that encodes for a Peptidoglycan recognition protein (PGRPLC) and is involved in the Imd pathway (Meister et al., 2009). This protein has been reported to be implicated in anti-*Plasmodium* responses at the level of the mosquito midgut, but without information about the functionality of their isoforms (Sreenivasamurthy et al., 2013). This gene consists of six isoforms corresponding to three different variants. PGRPLC gene produces three primary protein isoforms each with a distinct PGRP domain and an optional 75-nucleotide cassette located at the 3’ end of the shared exon 3. Consequently, each isoform is present in two forms, the longer version is 25 amino acids longer than the shorter version. AGAP005203 has 2 isoforms differentially expressed in our data all of them corresponding to the shorter version of the variants. RA form has been reported to be overexpressed during midgut infection and has been proposed to be involved in the mosquito immune response against *Plasmodium* (Meister et al., 2009). On the other side, the RE isoform, was overexpressed in salivary glands, and although its function in this tissue is still unknown. Previous work reported that the RD and RE isoforms are positively regulated in response to stress and antimicrobial responses (Lin et al., 2007, Chen, Ling and Weng, 2009, Meister et al., 2009).

Chromatin dynamics and transcriptional regulation are intimately related, but its role in the regulation of alternative splicing remains little known. There are very few studies that investigated how epigenetic mechanisms such as histone modifications or nucleosome positioning, would contribute to changes in the AS pattern. This is despite the functional consequences that AS events can have during development and disease (Simon et al., 2014, Petrova, Song, Consortium, NordströmNordstr, et al., 2022). Previous evidence in model organisms reported local changes in nucleosome positioning histone modifications and chromatin accessibility near the exon that undergoes splicing (Naftelberg et al., n.d., Chodavarapu et al., 2010, Li et al., 2020b).

Here, we proposed a model for *A. gambiae* to describe the association between isoform expression and/or usage and chromatin structure dynamics (Fig. 4). For differentially expressed isoforms we expected increased promoter accessibility in the tissue where the isoform was more expressed (Li et al., 2020b, Ruiz, Ranford-Cartwright and Gómez-Díaz, 2021). For the differentially used isoforms, we expected accessibility changes associates to the differential use, but localized at or nearby AS regulatory regions (splicing sites). In agreement with this model, our results indicate that there is a relationship between the differential expression and the use of isoforms with changes in chromatin accessibility enrichment suggesting that chromatin structure would play a role in AS-mediated gene expression regulation in *A. gambiae*. However, this relationship was only significant in the region of the isoform body. This is supported by previous studies in mammals and plants, reporting higher chromatin accessibility coinciding with the region of the intron retention (Ullah et al., 2018, Petrova, Song, Consortium, Nordström, et al., 2022).

**Fig 4.**
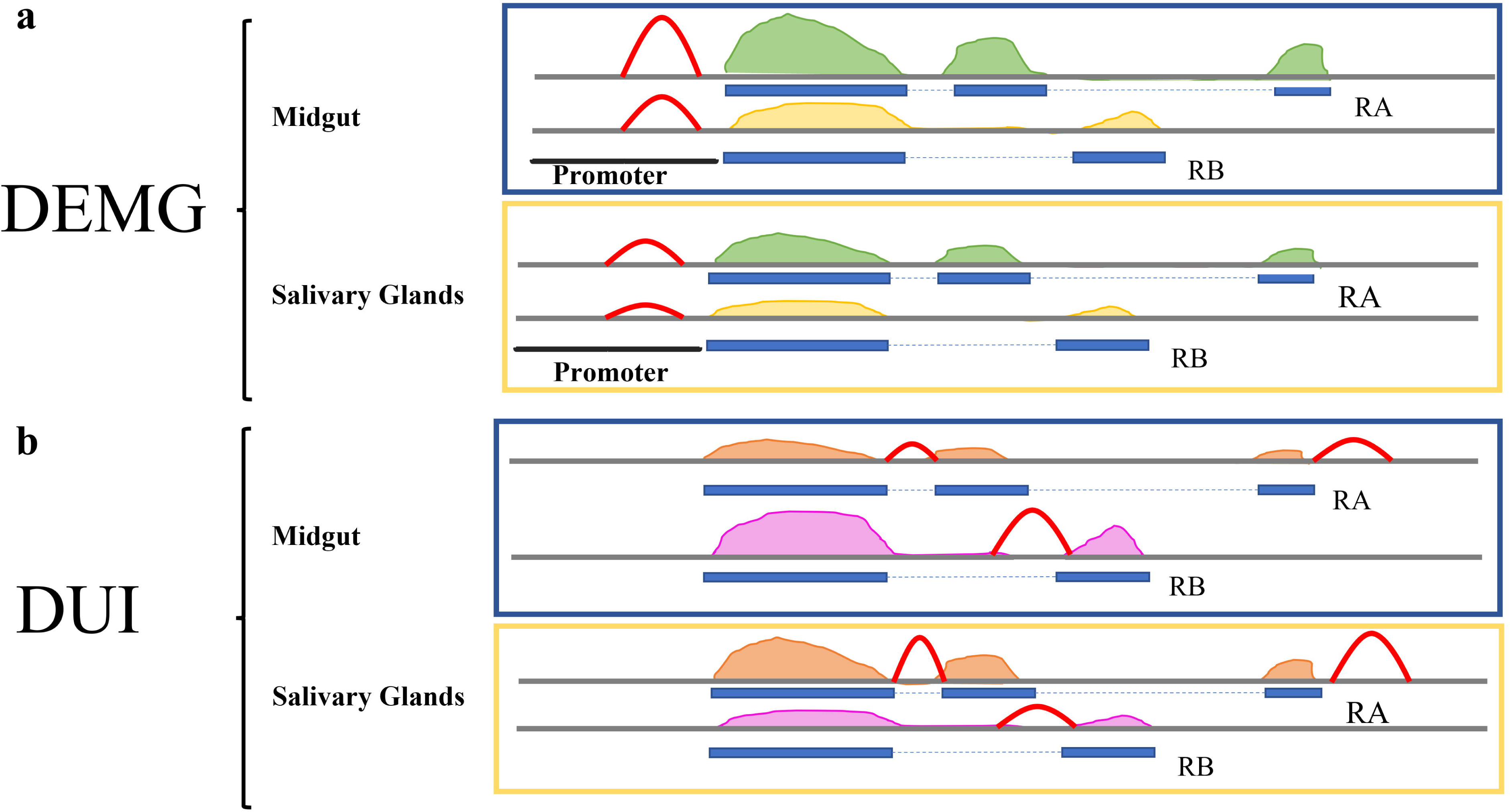
Model for the relationship between isoform expression, isoform usage and chromatin accessibility. The ATAC-seq peaks are shown in red. **a**, For the DEMG group the expectation is that the accessibility of the promoter will increase in the tissue where the expression of the isoform is higher i.e, The RA isoform in midguts display a higher ATAC-seq peak. For the RB isoform, the expression is higher in salivary glands the accessibility peak is higher in this tissue; **b**, For the DUI group, we expect to find chromatin accessibility changes near the splicing sites coinciding with the different usage of the isoform/s. The ATAC-seq peaks are around the differentially exons and are higher in the tissues where the isoforms are more used. The RA isoform is less expressed in midguts and becomes more expressed in salivary glands, displaying a higher ATAC-seq peak.

In addition, changes in accessibility along the isoform body have also been shown to alter the speed of RNA polymerase II (RNA pol II) (Saldi et al., 2016) or, alternatively, impact the recruitment of splicing factors through direct or indirect recognition of the different chromatin marks (Schor, Gómez Acuña and Kornblihtt, 2013). Indeed, an increase in chromatin accessibility in histone-depleted genes was shown to alter the elongation rate of Pol II, resulting in several transcripts with increased intron retention and changes in alternative splicing (Jimeno-González et al., 2015).

For differentially used isoforms, we already predicted that changes at the level of splicing sites would be more important. Our results confirm the existence of differential accessibility peaks at splicing sites for around 54% of the isoforms. What we cannot predict from our results, however, are the precise mechanism and the functional consequences of this change, that is, whether a local increase in chromatin accessibility at the splicing site would result in one or another alternatively spliced isoform due the selective binding of splicing repressor versus an inducer. Indeed, according to a recent study, the type of event and the choice of a particular isoform would be multifactorial and depend on a trade-off between chromatin accessibility at regulatory sequences (such as splicing sites), and variable RNA pol II elongation rates (Pai and Luca, 2019).

Motif analysis can give some insights on the underlying mechanisms of AS regulation. Looking at the *de novo* motifs identified in the differentially expressed and used isoforms, we found overrepresented motifs that resemble several transcription and slicing factors of *Drosophila* such as snail (sna), grainy head (grh) and chromatin-linked adaptor for MSL proteins (CLAMP). In the case of CLAMP, this is a chromatin remodeling protein with many different functions in regulating RNA Pol II dynamics, chromatin architecture, and alternative splicing (Schuettengruber et al., 2009) (Urban et al., 2017). It has been described for example, that the interaction of CLAMP with the proteins Squid (Sqd) and Syncrip (Syp), members of the heterogeneous nuclear ribonucleoproteins (hnRNPs), has a role in the regulation of the AS in *Drosophila* (Blanchette et al., 2009).

Another aspect that we approached in this study is the relationship between the enhancer function, based on histone PTMs and accessibility patterns, and AS events in *A. gambiae*. Previous evidence suggests that enhancers could mediate isoform-specific expression by regulating the production and binding of antisense transcripts, as well as by regulating messenger RNA 3′ end processing (Onodera et al., 2012, Shiau et al., 2021, Kwon et al., 2022). In such cases, enhancers were expected to be located intragenically, near the exons or introns that undergo AS (Lam and Hertel, 2002, Matlin, Clark and Smith, 2005).

When we look at the presence of enhancers in our AS susceptible isoforms (DEMG and DUI), we discovered that approximately 19-23% of differentially expressed and used isoforms have an enhancer annotated, and approximately 18-20% of these are active in our study, i.e. accessible. This observation suggests a relationship of the enhancers with the expression and use of the isoforms. Indeed, we also report that 20% of the enhancers active in differentially expressed group and 28% in differentially used isoforms correspond to differentially accessible enhancers, i.e. they have differential activation status, between tissues. Furthermore, most of our accessible enhancers (64.6% of the total of both groups) appear to be intragenic. Although their function in alternative splicing still needs to be experimentally confirmed, the fact that their activation status changes according to changes in the expression and use of the isoforms to which they are annotated, suggests a possible role in AS-mediated isoform expression regulation.

One example of a AS gene with an associated enhancer that is differentially accessible between conditions is the immune gene AGAP005234. In *A. gambiae* the function of this gene is associated the production of reactive oxygen species (ROS), that have been implicated in the immune response to pathogens (Marikovsky et al., 2003), including a *Plasmodium* infection (Molina-Cruz et al., 2008). Our data shows that the RB isoform is more used and expressed in salivary glands, whereas the RA isoform is only used in midguts. There is an enhancer annotated to the AGAP005234 gene that is more accessible in salivary glands, suggesting a possible regulatory function of this enhancer in determining the choice of which of the isoforms of this gene is used (RA or RB) or differentially expressed (RB).

To conclude, this is, to our knowledge, the first study that analyze comprehensively alternative splicing and its regulation in *A. gambiae*. We report genome-wide alterations in isoform expression and usage between different tissues and, less so, linked to the infection status. Also, the study of the relationship between AS with chromatin, points to the importance of chromatin structure dynamics at splicing sites in tissue-specific gene expression regulation. Finally, the identification of active enhancers associated to differentially expressed and used isoforms suggests a possible role of these regulatory elements on the splicing process. Our study lays the foundation for further research on the mechanisms of alternative splicing in *A. gambiae*, including the functional characterization of regulatory elements like enhancers and splicing factors as potential candidates for vector control.

## 4 Experimental Procedures

### 4.1 External data

External RNA-seq data analysed in this study was downloaded from GEO accession number GSE120076 and corresponds to samples infected and non-infected in midguts (Ruiz et al., 2019). Unpublished RNA-seq data corresponds to samples of salivary glands infected and non-infected from the same study (Ruiz et al., 2019), and has been assigned to GEO accession number GSE243215. Previously published ATAC-seq data for infected tissues, both in midgut and salivary glands, was downloaded from GEO accession number GSE152924 (Ruiz, Ranford-Cartwright and Gómez-Díaz, 2021).

### 4.2 Genes susceptible of alternative splicing and experimental design

To study the role of Alternative Splicing in *A. gambiae* we considered genes that are susceptible to be regulated by this mechanism. Only 1,263 genes in the *A. gambiae* genome (13,094 genes) have more than one transcript and could be classified as multisoform genes.

We used 12 RNA-seq and 4 ATAC-seq datasets corresponding to 8 *A. gambiae* infected and 4 non-infected (control) samples from two mosquito tissues (midguts and salivary glands). Midgut datasets correspond to published data from two previous studies(Ruiz et al., 2019, Ruiz, Ranford-Cartwright and Gómez-Díaz, 2021). And there is additional data from two samples (infected and non-infected salivary glands) that are new to this study. Samples are distributed in three different comparisons: infection *vs*. control in midguts and salivary glands, and infected midguts *vs*. infected salivary glands, having each of them 2 replicates per condition.

### 4.3 RNA-seq data analysis

Quality control of RNA-seq data was performed with FastQC v0.11.9 (Babraham Bioinformatics -FastQC A Quality Control tool for High Throughput Sequence Data, 2022). A replicability test based on correlation analysis and principal component analysis (PCA), support the use of the different samples as replicates (Supplementary figure 1).

We used SortMeRNA v4.2.0 (Kopylova, Noé and Touzet, 2012) to determine and eliminate possible ribosomal RNA contamination. Filtered reads were mapped to the reference genome (Anopheles gambiae PEST genome, version P4.12) available in VectorBase (release 53) (Giraldo-Calderón et al., 2015), using Salmon v.0.11.2 (Patro et al., 2017). This software is specifically used for the alignment and quantification of isoforms (Zhang et al., 2016) and was run using the standard parameters *-l A --validateMappings --go 4 --mismatchSeedSkip 5*. For this analysis we applied a conservative filter, only isoforms with a count value greater than 10 were considered as being expressed. A total number of 1,561 (Inf MG vs. Inf SG comparison), 1,394 (Inf vs. Ctrl MG), 1,679 (Inf vs. Ctrl SG) isoforms passed this threshold and were included in all subsequent analysis.

### 4.4 AS mechanisms and differential analyses

The counts generated by Salmon were used to perform various analyses. First, to study the most prevailing alternative splicing mechanisms in each condition we used the R package IsoformSwitchAnalyceR v1.12.0 (Vitting-Seerup, Sandelin and Berger, 2019). We applied the function *analyzeAlternativeSplicing* with the argument *onlySwitchingGenes=F*. We then compared the frequencies of the AS events between conditions (tissues and infection status) to determine how are they altered.

The differential expression analysis was performed using the R package (v4.0.2) DESeq2 v1.30.0, (Love, Huber and Anders, 2014). Taking multisoform genes, we considered differentially expressed isoforms (DEMG), those that have differential expression between conditions (the isoform not the gene). To be differentially expressed, isoforms must have a significant P-value of less than 0.05 and a LogFoldChange greater than 1 or less than -1 (corresponding to a 10% change). In the comparison between Inf MG vs. Inf SG a LogFC < 0 means overexpression in MG, whereas a LogFC > 0 means overexpression in SG. In the comparison between Inf vs. Control a LogFC > 0 means more expression in the infected tissue.

The study of the differential usage of the isoforms was performed using IsoformSwitchAnalyceR v1.12.0 (Vitting-Seerup, Sandelin and Berger, 2019). In this case we used the parameter, difference in isoform fraction (dIF) to measure the magnitude of change. The isoform fraction (IF) is a measurement that determines the proportion of the total expression of a parent gene that comes from a particular isoform (isoform_exp / gene_exp). Isoforms significantly differentially used (DUI) were those with a significancy P-value below 0.05. In the comparison between Inf MG vs. Inf SG a dIF < 0 means more used in MG, and dIF > 0 means more used in SG. In the comparison between Inf vs. Control a dIF < 0 means more usage in the infected tissue.

We performed Gene Ontology (GO) terms overrepresentation tests for the sets of differentially expressed or used multisoform genes between Infected midguts *vs.* Infected salivary glands using VectorBase Overrepresentation Test and ImmunoDB database to know which isoforms are immunologically relevant (Giraldo-Calderón et al., 2015, EZlab | Computational Evolutionary Genomics group, 2022). We chose the p-value as a measure of the likelihood that a certain GO term appear among the genes in your results more often than it appears in the set of all genes for that organism (background) and we applied a threshold of p-value < 0.05.

### 4.5 ATAC-seq data analyses

For the analysis of ATAC-seq data in relation to AS, we used processed data from a recent article previously published by our group (Ruiz, Ranford-Cartwright and Gómez-Díaz, 2021) the nucleosome free reads, the ATAC peaks resulting from the analysis with MACS2 (Zhang et al., 2008a) and the differentially accessible peaks resulting from the analysis with Diffbind (Stark R. and Brown G., 2011).

In brief, raw reads were trimmed 20 bases from the 3′ end of each read (−3 20) and aligned to the *A. gambiae* PEST genome with Bowtie2 (Langmead and Salzberg, 2012) (v2.4.1) using default parameters, except for –no-unal –no-mixed -X 2000. We then applied a MAPQ score threshold of 10 and sorted and deduplicated the reads using SAMtools (Li et al., 2009) (v1.10). To adjust the known bias and ensure the mapping of Tn5 cutting sites, we shifted aligned reads +4 bp for + strands and −5 bp for − strands with ATACSeqQC (Ou et al., 2018) (v1.10). We removed not properly paired reads and extracted nucleosome-free fragments with a size threshold of 130 bp (SAMtools). We performed peak-calling on nucleosome-free reads with MACS2 (Zhang et al., 2008b) (v.2.1.2) using the callpeak module and the following parameters: -f BAMPE -g 273109044 -q 0.01 -B –keep-dup all –nomodel –nolambda. We refer to ATAC-seq peaks as Tn5 hypersensitive sites (THSs). To identify THSs unique or common across samples, we used BEDTools intersect (Quinlan and Hall, 2010) requiring a minimum overlap of 51% (- f/-F 0.51). We annotated THSs to genomic features combining HOMER (Heinz et al., 2010) (v4.11).

From the nucleosome free reads, we quantified the ATAC-seq nucleosome-free signal enrichment at our genomic regions of interest (promoter, isoform body) using BEDTools v2.30.0 (Quinlan and Hall, 2010) with the parameter *intersect* and the option ***-c***. Read counts were normalized (RPKM).

We used the DiffBind R package (Stark R. and Brown G., 2011) (v2.14) to assess differential chromatin accessibility at given locations between P. falciparum-infected mosquito MGs and SGs. As input for DiffBind, we used the ATAC-seq nucleosome-free reads and the THSs. Infection 1 and Infection 2 were used as biological replicates.

### 4.6 Relationship between gene expression and chromatin accessibility

#### 4.6.1 Model and predictions

The relationship between the gene expression and chromatin structure has been examined in a few model organisms (Luco et al., 2011). These previous studies showed that various epigenetic mechanisms such as nucleosome positioning, post-transcriptional modifications (Luco et al., 2011) or the modification of the chromatin accessibility (Li et al., 2020b, Nussinov et al., 2021), alter or correlate with AS dynamics. In *A. gambiae*, an earlier work investigated the relationship between chromatin accessibility and gene expression regulation at gene level (Ruiz, Ranford-Cartwright and Gómez-Díaz, 2021), but here we focused on gene regulation occurring at the isoform level via alternative splicing.

We made some predictions on how chromatin structure may be altered when comparing different mosquito tissues in which AS would play a role in gene expression regulation (see previous section), and the pattern expected considering DEMG and DUI genes. Our expectation was to find a positive relationship between chromatin accessibility differences and the differential expression and/or usage of the isoforms (Fig 4).

For the differentially expressed group, we expected to find an increased accessibility in the promoter of one or more isoforms in the tissue where the expression of the isoform/s is higher (Fig 4a). In the differentially used isoforms group, we expected to find changes in chromatin accessibility at regulatory regions coinciding or near alternative splicing sites in the isoform more used in each tissue, as is shown in Fig 4b. For example, for an exon skipping event we expected more accessibility in the surroundings of the exon to be added to the transcript. In our data the alternative splicing event more used is the ES and the majority of the MACS2 peaks were in the isoform body. Alternatively, this association could also be evident in the promoter of isoforms differentially used for AS events such as ES, ATTS and IR. (Li et al., 2020b).

#### 4.6.2 Data analysis

To test the possible relationship between differential expression of isoforms and chromatin accessibility, we discarded genes that overlap by more than 25% with other genes. We used the Spearman’s correlation test from R v4.2.0 to analyze the possible association between isoform expression and chromatin accessibility at various regions where the AS mechanisms would operate: The isoform promoter, which comprises 1Kbp from the TSS (end 5’ UTR), and the isoform body including all exons and introns. For those isoforms that lack a TSS annotated, we considered the ATG as the reference point (supplementary figure 2).

We used a R script to classify the multisoform genes into three groups - high, medium, and low - depending on the expression level of the isoforms. The threshold values for this classification were determined by dividing the signal into three quantile groups based on their means, using the Hmisc::cut2 R function. We then used the Spearman’s correlation test to compare the isoform expression and chromatin accessibility for each group separately. The software *ngsplot* was used to visualize as Average profile plots the chromatin accessibility (nucleosome free region signal) at the TSS in the different groups of isoforms classified by their expression level (high, medium, low) (Shen et al., 2014) (Supplementary figure 3).

In the case of differentially used isoforms, we analyse the association between isoform usage and chromatin accessibility focusing on alternative splicing sites. To do this, we first searched for significantly differential peaks between conditions (previously obtained by DiffBind) that coincide with *A. gambiae* known splicing sites (SS) adding a small distance to see what is happening around these sites. We set +/- 100bp being restrictive, and subsequently we counted how many of those peaks annotate to isoforms classified as differentially used. On those peaks, we then we performed a motif analysis using HOMER software v4.11 (Homer Software and Data Download, 2022). We used *findMotifGenome.pl* to determine known and *de novo* motifs overrepresented in our sequences, and then used *annotatePeaks.pl* to link the motifs back to the differentially used isoforms and the splicing sites. Only motifs with significance p-values less than 1e-25 were considered.

### 4.7 Enhancers associated to AS in *A. gambiae*

Based on the integrative analysis of the ATAC-seq and RNA-seq data, we used the set of *A. gambiae* enhancers characterized in a previous study (Ruiz, Ranford-Cartwright and Gómez-Díaz, 2021), to interrogate whether these regulatory sequences could be involved in AS events. The list included 1,402 *A. gambiae* enhancers predicted by homology with *Drosophila* enhancers and, 2,866 *Anopheles* enhancers identified either computationally or experimentally by different studies (Ruiz, Ranford-Cartwright and Gómez-Díaz, 2021). We selected only the enhancers that appeared to be active in at least one of our conditions, that is, enhancers that have a MACS2 chromatin accessibility peak and appear enriched in H3K27ac. As a result, we obtained a list of 811 active enhancers which we overlapped with our differentially expressed and used isoforms using BEDtools *intersect*. We also classified these enhancers as intragenic or intergenic using BEDtools *intersect*. Finally, we quantified which portion of the active enhancers that annotated to differentially expressed and used isoforms, display a change in accessibility; that is, are differentially accessible between tissues coinciding with the change in the expression of the isoform/s.

## Abbreviations

DEMG: Differential isoform expression of multisoform genes;
DUI: Differential isoform usage of multisoform genes; SS, splicing sites.

## Acknowledgements

We are grateful to Dr Lisa Ranford-Cartwright and Dr José L. Ruiz for sharing data. We thank the bioinformatics unit of the IPBLN-CSIC for computing facilities and advice on the analysis.

## Tables

**Table 1.**
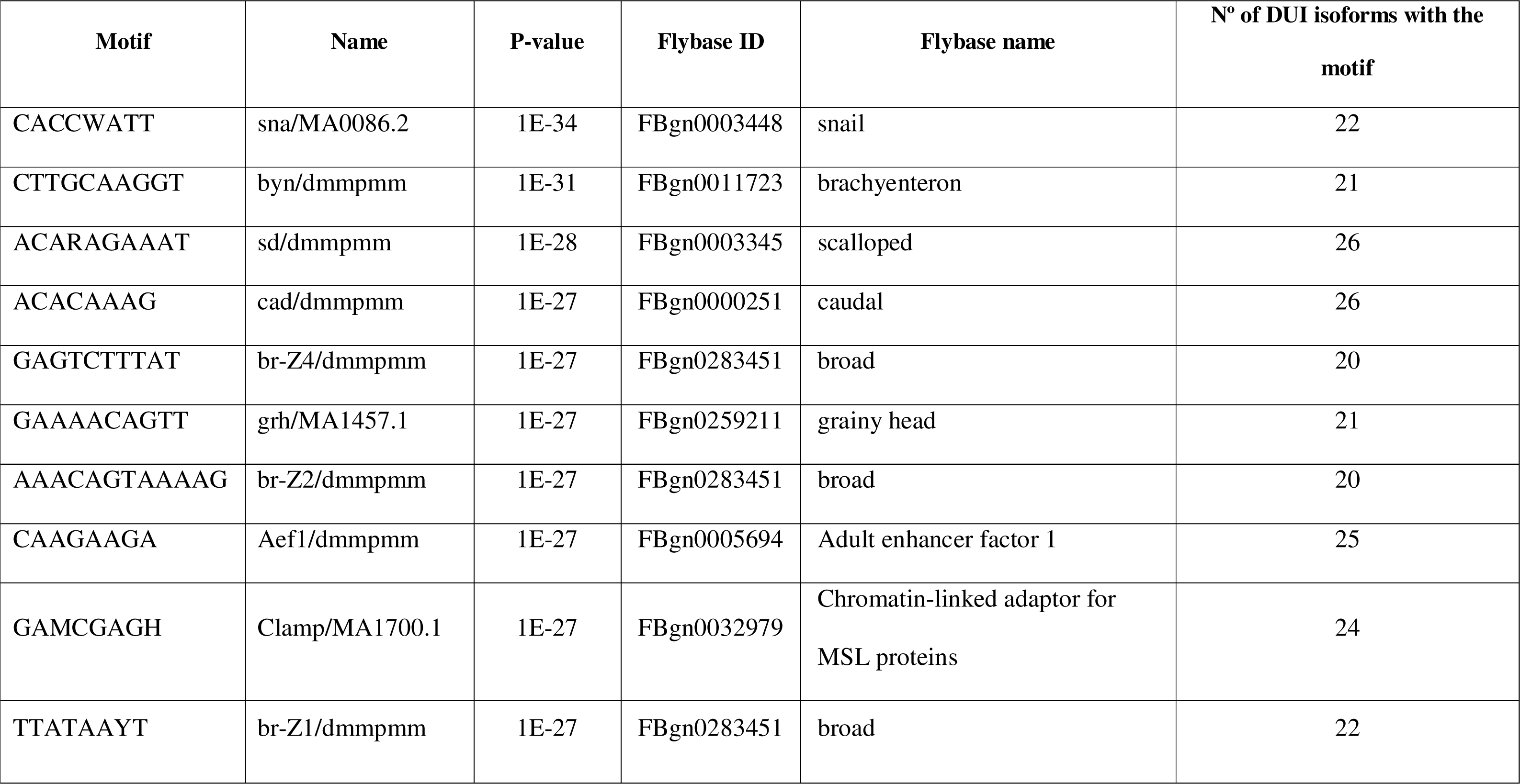
Ten first “de novo” motifs identified by Homer that appear overrepresented in the splicing sites of differentially expressed and used isoforms that display differential accessibility between tissues. Additional information in Supplementary table S6.

