## Supplementary table legends and figures for "Alternative splicing and its regulation in the malaria vector *Anopheles gambiae*"

### Supplementary material

#### Supplementary tables legends

**Supplementary Table S1.** Differential isoform expression analysis.

Differentially Expressed isoforms corresponding to multisoform genes (DEMGs) for each experimental comparison. Columns include the isoform ID, DESeq2 output (Isoform, LogFoldChange, P-value…), tissue or infection status comparison and the condition where each isoform is more expressed.

**Supplementary Table S2.** Differential isoform usage analysis.

Differentially used isoforms corresponding to multisoform genes (DUIs) for each experimental comparison. Columns include the isoform ID, IsoformSwichAnalyzer output (Isoform, dIF, P-value, IF of each condition…), tissue or infection status comparison and the condition where each isoform is more used.

**Supplementary Table S3.** Gene ontology terms overrepresentation analysis.

Gene ontology (GO) terms overrepresentation analysis for the set of DEMGs and DUIs. Results include the biological processes (BP) GO terms.

**Supplementary Table S4.**  DEMG and DUI isoforms with vector-pathogen interactions related functions.

DEMG or DUI isoforms matching genes previously described by Sreenivasamurthy et al. (Sreenivasamurthy et al., 2013) with a known function in vector pathogen interactions.

**Supplementary Table S5. RNA-seq and ATAC-seq correlation analysis in the DEMG group**.

**(A)** Results of the correlation between gene expression and chromatin accessibility for DEMG isoforms overexpressed in midguts or salivary glands considering the promoter region or the isoform body. The rho and p-values are included. **(B)** Results of the correlation between gene expression and chromatin accessibility dividing isoforms by the level of expression (high, medium y low) for DEMG isoforms overexpressed in midguts or salivary glands. The rho and p-values are included.

**Supplementary Table S6.** Summary of the results of the motif analysis by Homer on the set of DiffBind peaks located at splicing sites for the DUI isoforms.

**(A)** List of known motifs including their p-value, % of sequences containing the motif and the annotation to *A. gambiae* DUI isoforms **(B)** List of 10 most significant *de novo* motifs ranked according to their p-value, the *Drosophila* TF showing similarity to the motif (including FlyBase ID and function), and annotation to DUI *A. gambiae* isoforms.

**Supplementary Table S7.** Active enhancers with a possible function in the regulation of AS in *A. gambiae*.

List of enhancers annotated to DEMG (A) and DUI (B) isoforms. The table contain the information of: type of gene, enhancer ID, type of enhancer, gene regulated, differentially accessible.

#### Supplementary Figures

**Supplementary Figure S1.**

Principal Component Analysis (PCA) showing the percentage of the variance being explained by the first 2 components (PC). Samples are grouped by condition (infection or control) rather than by experimental infection. The heatmap represents the degree of correlation (similarity) between samples. Samples from different experimental infections show higher correlation for the same condition than between different conditions. (A) Comparison Inf *vs*. Control Midguts (B) Comparison Infection *vs*. Control Salivary glands (C) Infection Midguts *vs.* Infection Salivary Glands.

**Supplementary Figure S2.**

Density plot showing the position of Tn5 hypersensitive sites (THSs) with respect to the TSSs (or the ATG for genes without annotated 5′ UTRs). Higher densities of THSs occur within 1 Kb upstream the TSSs or ATG sites. The dashed lines indicate the putative promoter region located 1 Kb upstream.

**Supplementary Figure S3.**

Profile plots (up) and violin plots (down) showing changes in ATAC-seq nucleosome-free signal enrichment at each tissue with respect to the TSSs (±1 Kb). Genes are divided into groups and ranked by their mRNA levels (high, medium or low). In the violin plots the width of the graph takes into account the density of repeated values in the interval. The mean values are marked with a black dot.

**Supplementary Figure S1.**

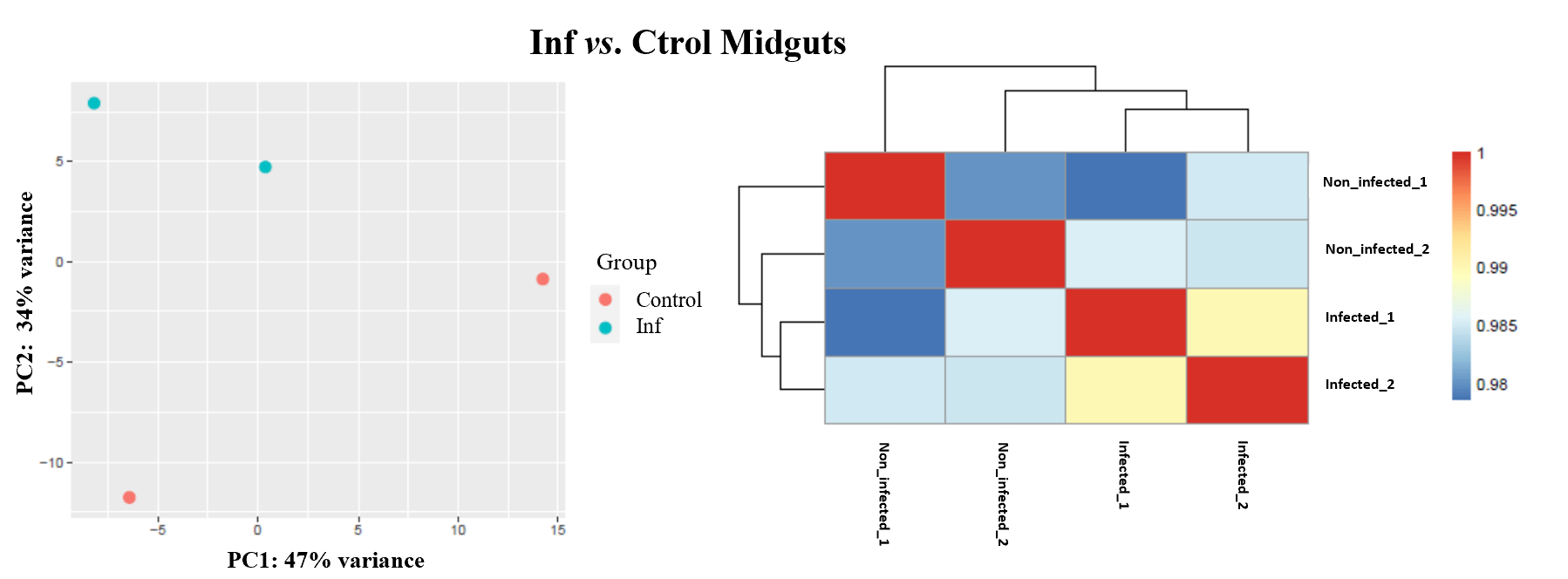

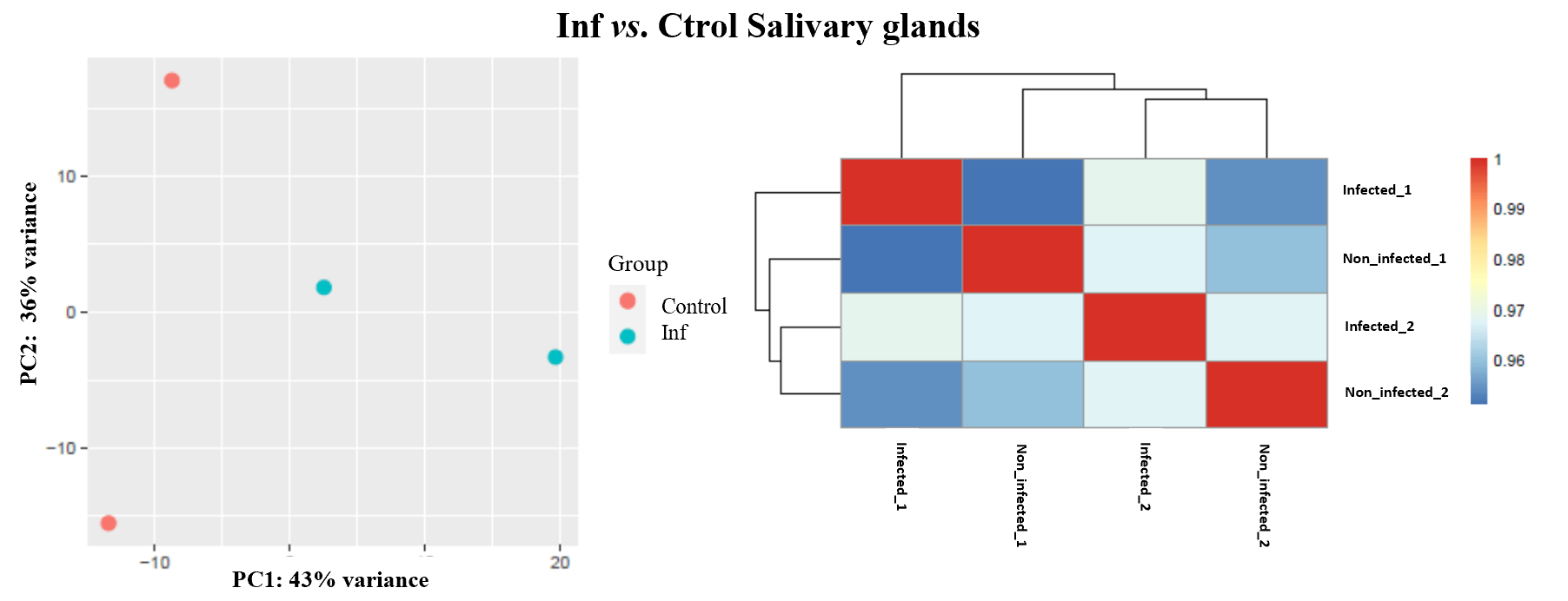

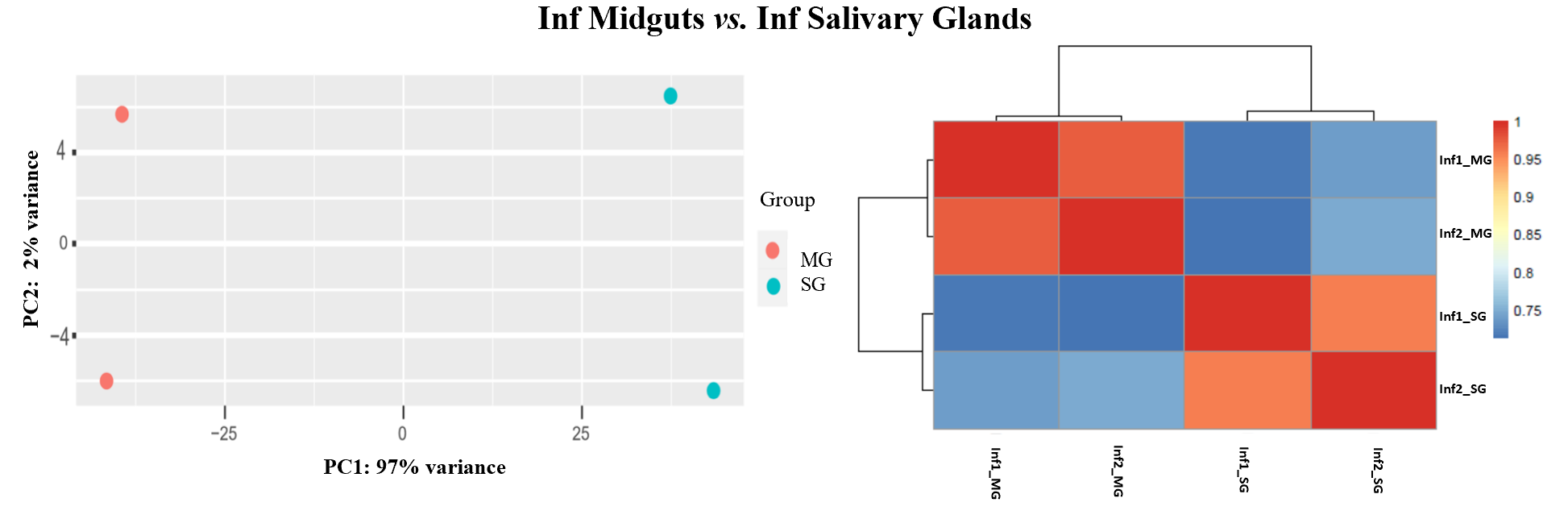

**Supplementary Figure S2.**

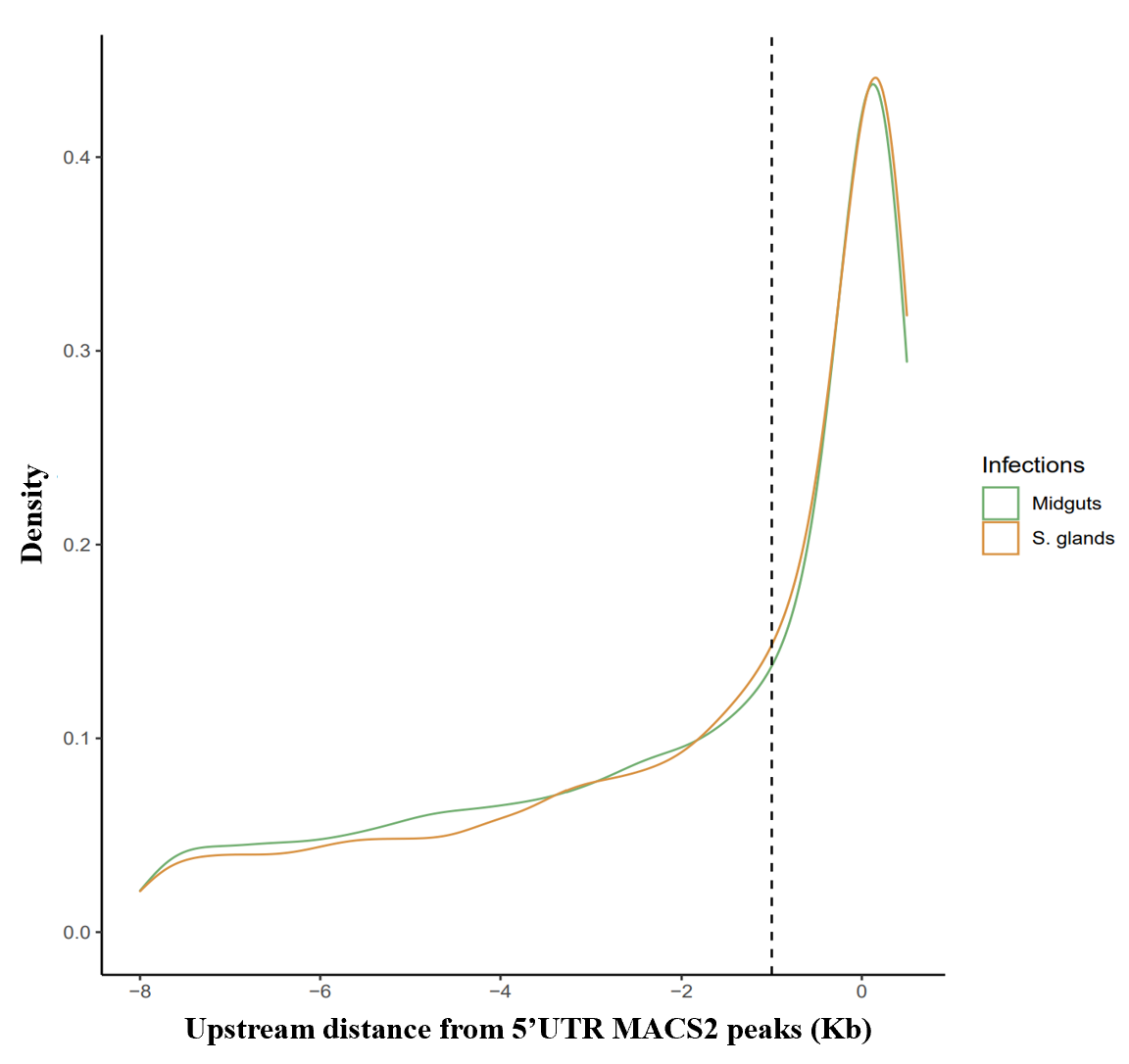

**Supplementary Figure S3.**

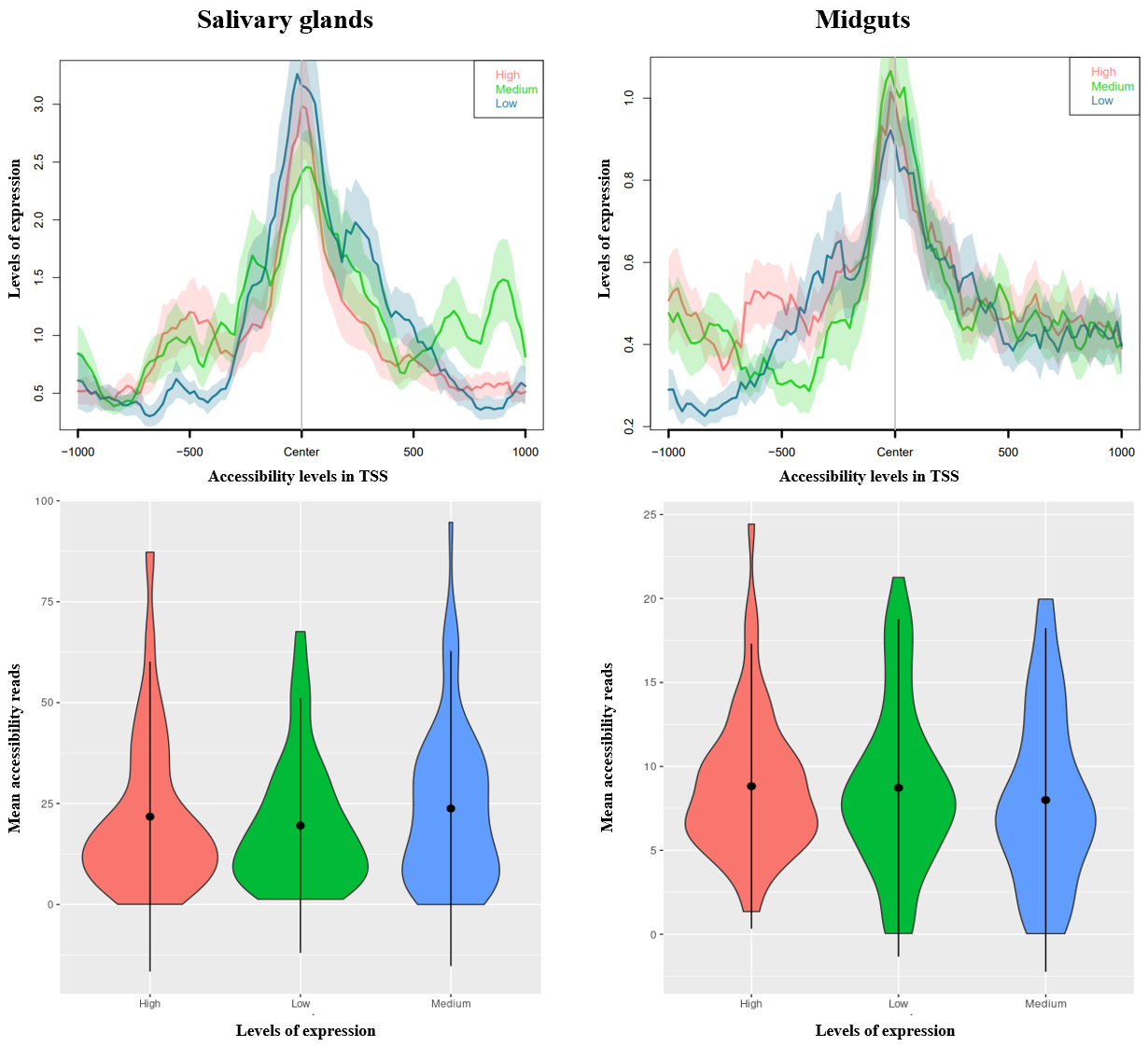
